# Big and Small Cerebral Asymmetries

**DOI:** 10.1101/269571

**Authors:** Mark A. Eckert, Kenneth I. Vaden, Dyslexia Data Consortium

## Abstract

Deformation-based asymmetries replicate previously observed grey matter asymmetries and characterize white matter asymmetries.

Increased sensitivity to structural asymmetries in some brain regions depends on smaller-scale normalization or deformation parameters.

Tuning deformation parameters can provide more precise asymmetry measures for understanding the mechanisms and functional significance of cerebral asymmetries.

---

The functional significance of hemispheric asymmetries in brain structure remains an intriguing and largely unexplained characteristic of brain organization (Güntürkün & Ocklenburg 2017; Toga & Thompson 2003). Frontal and occipital petalia, leftward planum temporale asymmetry, rightward superior temporal sulcus asymmetry, and leftward anterior insula asymmetries are consistently observed asymmetries in voxel-based morphometry studies in children (Eckert et al., 2008) and adults (Good et al., 2001; Watkins et al., 2001). Here we demonstrate that these asymmetries can be observed using deformation information that specifies how to normalize the images to a symmetrical template. We also show that identifying some structural asymmetries, including the genu of the corpus callosum for example, depends on the spatial scale of normalization.

The T1-weighted images from 306 normal participants (115 female; mean age 9.12 +/− 1.77 years) were contributed from 6 research sites and shared for a retrospective multi-site study with Institutional Review Board approvals. The native space T1-weighted images were rigidly aligned and left-right flipped to create a study-specific symmetrical template using diffeomorphic normalization (ANTS; Avants et al., 2011). This approach differs from typical voxel-based analyses where images are first segmented before normalization to a symmetrical template and then modulated to adjust grey matter voxel values to account for volumetric displacement to the template. Specifically, we used an ANTS Greedy SyN normalization with three steps of increasingly smaller resolution to examine the relative scale of different brain asymmetries when native space T1 images were aligned to the symmetrical T1 template [Parameters for normalization steps 1-3: Cross-correlation metric (mm radius = 4.00, 2.00, 1.00); SyN (1.00, 0.75., 0.50); Gaussian regularization (3.00, 2.00, 1.00); iterations (50, 50, 40)]. The spatially normalized log Jacobian fields were obtained from Step 1-3 deformation fields, smoothed with a Gaussian kernel (8 mm) as in previous structural asymmetry studies, and left-right subtracted to create log-scale hemispheric asymmetries. One-sample t-tests, adjusting for potential research site effects with voxel-level family-wise error correction for multiple comparisons (FWE, p < 0.05), were performed using SPM12 to identify significant asymmetries in volumetric displacement relative to the symmetrical template.

Figure 1A shows hemispheric structural asymmetries where grey matter asymmetries have been observed when Jacobian modulation was used, including leftward planum temporale asymmetry, rightward frontal and leftward occipital petalias, leftward anterior insula asymmetry, and rightward temporal pole asymmetry (Eckert et al., 2008; Good et al., 2001; Watkins et al., 2001; and cortical volume: Maingault et al., 2016). Inspection of the Jacobian asymmetries at each normalization step revealed that structures like the medial planum temporale, anterior insula, and superior temporal right sulcus had asymmetries that required all three normalization steps to align these regions to the symmetrical template. For example, participants with relatively greater asymmetry at Step 1 also had greater asymmetry at subsequent steps (e.g., medial planum temporale asymmetry Step 1 * Step 3: r_(304)_ = 0.31, p = 2.8759E-8). This appeared to be due to cases with the largest brain volumes. Leftward planum asymmetry at each normalization step was significantly related to total intracranial volume [TICV^1^ (Step 1: r_(304)_ = 0.12, p = 0.04; Step 2: r_(304)_ = 0.20, p = 0.001; Step 3: r_(304)_ = 0.20, p = 0.001]. Unexpectedly, this TICV association with planum temporale asymmetry was driven by females for the first two normalization steps (Step 1: r_(148)_ = 0.17, p = 0.04; Step 2: (r_(154)_ = 0.21, p = 0.01) and by the males for the last normalization step (r_(154)_ = 0.28, p = 0.0007). There were no significant asymmetry differences between males and females or age effects.

These results indicate that some of the findings from voxel-based grey matter asymmetry studies can be attributed to hemispheric differences in gross volumetric displacement that is represented in the normalization deformation fields rather than specific structural differences in grey matter volume. This is most clearly seen in the frontal petalia results where the asymmetries were largely in white matter. Similar results were observed for the medial planum temporale where this asymmetry extended into white matter and the parietal operculum (Figure 1A).

Some of the structural asymmetries were observed only for the final and smallest spatial scale of normalization (Step 3). For example, a parasagittal region of the corpus callosum genu exhibited a significant asymmetry (Figure 1B) because of the final step of warping (Figure 1D). This asymmetry appeared to be due to both the anterior-to-posterior thickness of the parasagittal genu, which may relate to fractional anisotropy asymmetry in the parasagittal genu (Park et al., 2004), as well as a modest anterior to posterior shift in position compared to the right genu (Figure 1F,G). In contrast to the medial frontal asymmetry results described above, genu asymmetry at Step 3 was not significantly related to genu asymmetries at Step 1 (r_(304)_ = 0.11, p = 0.06), but was significantly related to genu asymmetry at Step 2 (r_(304)_ = 0.41, p = 4.7431E-14). The larger scale warping parameters were not sensitive to or deterministic for this smaller scale asymmetry, which is supported by the observation that genu asymmetry at Step 3 was not significantly related to TICV (females: r = 0.09, ns; males: r = 0.05, ns).

Asymmetry results specific to a smaller spatial scale of resolution highlight the potential to optimize normalization parameters to enhance sensitivity for brain structures of interest. The smaller scale normalization parameters also identified asymmetry in sulcal regions, such as the anterior superior temporal sulcus (Table 1), where there can be substantial variation in morphology. Together, the results suggest that locally specific and gross morphologic asymmetries, which we predict may be explained by mechanical forces (Ferreira & Vermot, 2017) and epigenetics (Schmitz et al., 2017), respectively, can be characterized to advance understanding about the functional significance of brain structure asymmetries.

**Figure 1.**
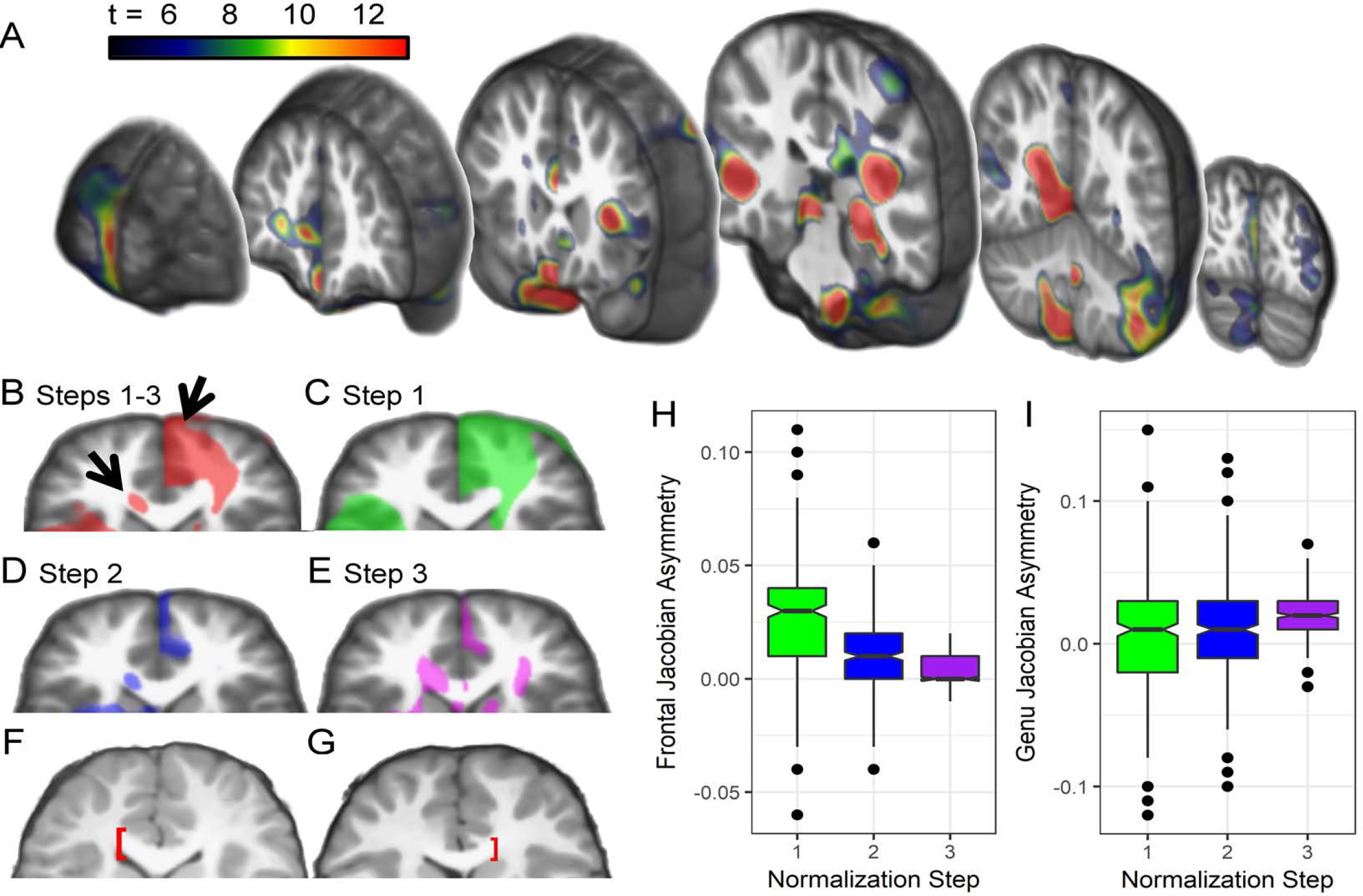
Deformation-based asymmetries. A. Consistent with voxel-based findings, there were significant leftward asymmetries in the medial planum temporale/Heschl’s gyrus, occipital pole, and anterior insula/inferior frontal gyrus asymmetries, as well as significant rightward asymmetries in the superior temporal sulcus, medial temporal lobe, and frontal pole (FWE p < 0.05). B. Arrows highlight rightward medial frontal and leftward parasagittal genu asymmetries that are shown for each normalization step in C-E (all FWE p < 0.05; axial plane of section). C. The medial frontal effect is largely due to the first warping step with the largest spatial resolution. D,E. The parasagittal genu asymmetry was most pronounced at the warping step with the finest spatial resolution. Note the relatively smaller extent of frontal asymmetry effects at the smaller scale of warping. F. 9.71 year-old male with leftward parasagittal genu asymmetry. G. 7.75 year-old male with rightward parasagittal genu asymmetry. F,G. Note the more anterior position of the genu in the hemisphere with the more pronounced leftward or rightward deformation asymmetry. H,I. Box-plots show that frontal and genu asymmetries were dependent on large volumetric displacements (Step 1) versus smaller scale displacement (Step 3), respectively (y-axes reflect the left-right subtraction of log Jacobian values). The 306 cases included 61 who were 18.69-28.05 years of age. These adults were included to replicate asymmetry findings in the younger sample and did exhibit the same spatial pattern of effects, including the genu and frontal petalia effects, although the asymmetry t-scores were lower for the adults due to the smaller number of cases. Because the results were consistent across ages and because of an absence of age effects (p < 0.05, FWE corrected), results for all 306 cases are presented.

**Table 1.**
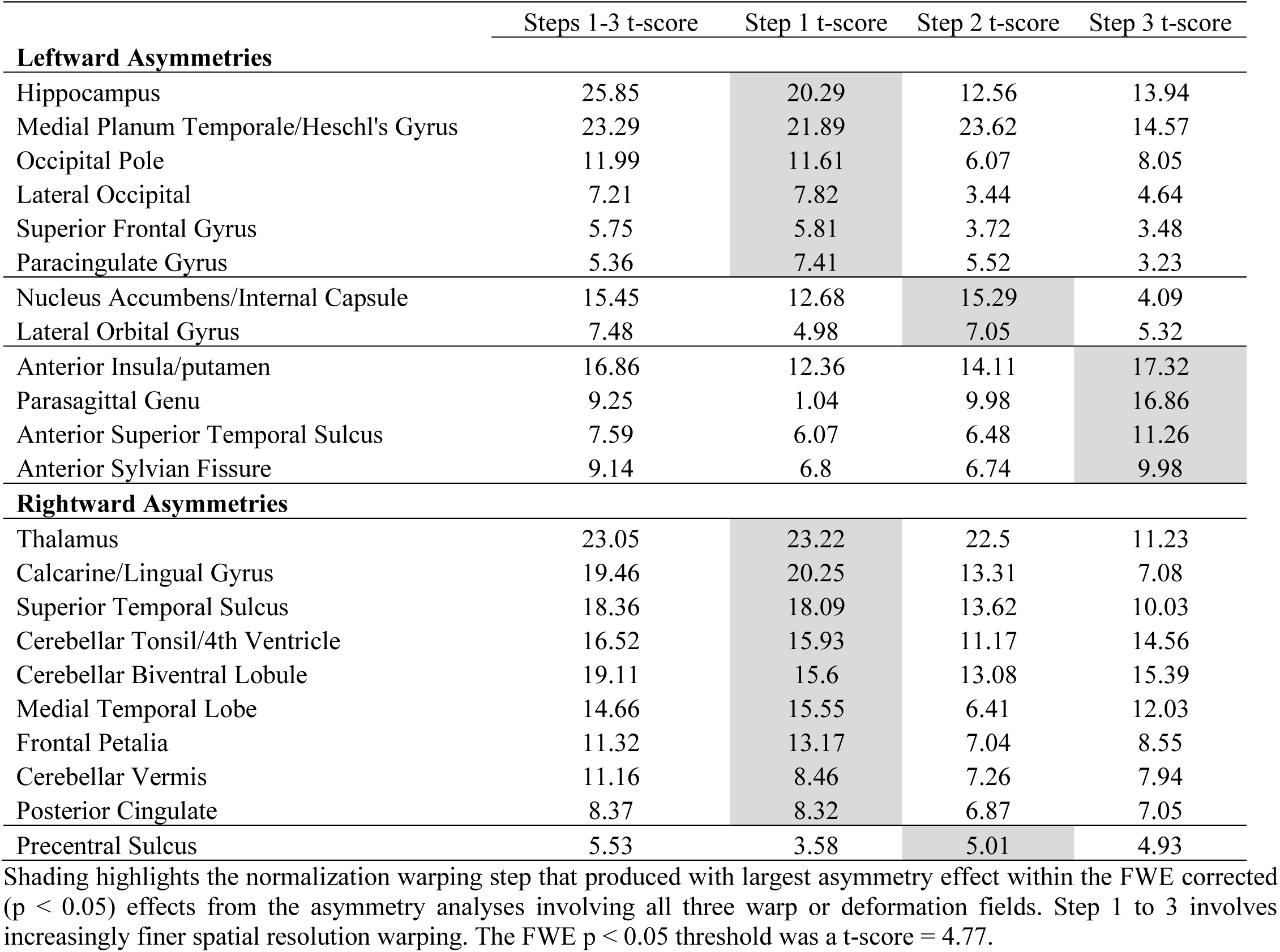
Cerebral asymmetries by the normalization step that contributed most to the asymmetry.

|  | Steps 1-3 t-score | Step 1 t-score | Step 2 t-score | Step 3 t-score |
| --- | --- | --- | --- | --- |
| **Leftward Asymmetries** |  |  |  |  |
| Hippocampus | 25.85 | 20.29 | 12.56 | 13.94 |
| Medial Planum Temporale/Heschl's Gyrus | 23.29 | 21.89 | 23.62 | 14.57 |
| Occipital Pole | 11.99 | 11.61 | 6.07 | 8.05 |
| Lateral Occipital | 7.21 | 7.82 | 3.44 | 4.64 |
| Superior Frontal Gyrus | 5.75 | 5.81 | 3.72 | 3.48 |
| Paracingulate Gyrus | 5.36 | 7.41 | 5.52 | 3.23 |
| Nucleus Accumbens/Internal Capsule | 15.45 | 12.68 | 15.29 | 4.09 |
| Lateral Orbital Gyrus | 7.48 | 4.98 | 7.05 | 5.32 |
| Anterior Insula/putamen | 16.86 | 12.36 | 14.11 | 17.32 |
| Parasagittal Genu | 9.25 | 1.04 | 9.98 | 16.86 |
| Anterior Superior Temporal Sulcus | 7.59 | 6.07 | 6.48 | 11.26 |
| Anterior Sylvian Fissure | 9.14 | 6.8 | 6.74 | 9.98 |
| **Rightward Asymmetries** |  |  |  |  |
| Thalamus | 23.05 | 23.22 | 22.5 | 11.23 |
| Calcarine/Lingual Gyrus | 19.46 | 20.25 | 13.31 | 7.08 |
| Superior Temporal Sulcus | 18.36 | 18.09 | 13.62 | 10.03 |
| Cerebellar Tonsil/4th Ventricle | 16.52 | 15.93 | 11.17 | 14.56 |
| Cerebellar Biventral Lobule | 19.11 | 15.6 | 13.08 | 15.39 |
| Medial Temporal Lobe | 14.66 | 15.55 | 6.41 | 12.03 |
| Frontal Petalia | 11.32 | 13.17 | 7.04 | 8.55 |
| Cerebellar Vermis | 11.16 | 8.46 | 7.26 | 7.94 |
| Posterior Cingulate | 8.37 | 8.32 | 6.87 | 7.05 |
| Precentral Sulcus | 5.53 | 3.58 | 5.01 | 4.93 |

Shading highlights the normalization warping step that produced with largest asymmetry effect within the FWE corrected ( $p < 0.05$ ) effects from the asymmetry analyses involving all three warp or deformation fields. Step 1 to 3 involves increasingly finer spatial resolution warping. The FWE $p < 0.05$ threshold was a t-score = 4.77.

## Acknowledgements

Members of the Dyslexia Data Consortium during preparation of this manuscript include C. Beaulieu, V. Berninger, X. Castellanos, C. Chiarello, T. Conway, L. Cutting, G. Dehaene-Lambertz, G. Eden, R. Frye, D. Giaschi, J. Gilger, F. Hoeft, M. Kibby, K. van Krigstein, M. Kronbichler, C. Leonard, M. Milham, T. Odegard, R. Poldrack, K. Pugh, T. Richards, N. Rollins, K. Schneider, J. Talcott, and B. Wandell. This work was supported by 5R01HD069374. This investigation was conducted in a facility constructed with support from Research Facilities Improvement Program (C06 RR14516) from the National Center for Research Resources, National Institutes of Health.

## Footnotes

1 SPM12 was used to segment the native space T1 images and calculate total intracranial volume based on the summed voxel probability values for grey matter, white matter, and CSF.

## Notes

The authors have no conflicts of interest.

